# Machine-learning to Improve Prediction of Mortality following Acute Myocardial Infarction: An Assessment in the NCDR-Chest Pain-Myocardial Infarction Registry

**DOI:** 10.1101/540369

**Authors:** Rohan Khera, Julian Haimovich, Nate Hurley, Robert McNamara, John A Spertus, Nihar Desai, Frederick A. Masoudi, Chenxi Huang, Sharon-Lise Normand, Bobak J. Mortazavi, Harlan M Krumholz

**Affiliations:** Division of Cardiology, University of Texas Southwestern Medical Center, Dallas, TX; Department of Internal Medicine, Massachusetts General Hospital, Boston, MA; Computer Science & Engineering, Texas A&M University, College Station, TX; Section of Cardiovascular Medicine, Department of Internal Medicine, Yale School of Medicine, New Haven, CT; Saint Luke’s Mid America Heart Institute, Kansas City, MO; Division of Cardiology, Department of Internal Medicine, University of Missouri-Kansas City, Kansas City, MO; Center for Outcomes Research and Evaluation, Yale-New Haven Hospital, New Haven, CT; Division of Cardiology, Department of Internal Medicine, University of Colorado Anschutz Medical Campus, Aurora CO; Department of Biostatistics, T.H. Chan School of Public Health, Harvard University, Boston, MA; Department of Health Care Policy, Harvard Medical School, Boston, MA; Department of Health Policy and Management, Yale School of Public Health, New Haven, CT

**Author notes:** Contributed equally as co-first authors (Rohan Khera, Julian Haimovich). **Corresponding Author:** Dr. Harlan M. Krumholz; 1 Church St., Suite 200, New Haven, CT 06510.

**Keywords:** Machine learning, AMI, risk prediction, outcomes research

## Abstract

**Introduction:** Accurate prediction of risk of death following acute myocardial infarction (AMI) can guide the triage of care services and shared decision-making. Contemporary machine-learning may improve risk-prediction by identifying complex relationships between predictors and outcomes.

**Methods and Results:** We studied 993,905 patients in the American College of Cardiology Chest Pain-MI Registry hospitalized with AMI (mean age 64 ± 13 years, 34% women) between January 2011 and December 2016. We developed and validated three machine learning models to predict in-hospital mortality and compared the performance characteristics with a logistic regression model. In an independent validation cohort, we compared logistic regression with lasso regularization (c-statistic, 0.891 [95% CI, 0.890-0.892]), gradient descent boosting (c-statistic, 0.902 [0.901-0.903]), and meta-classification that combined gradient descent boosting with a neural network (c-statistic, 0.904 [0.903-0.905]) with traditional logistic regression (c-statistic, 0.882 [0.881-0.883]). There were improvements in classification of individuals across the spectrum of patient risk with each of the three methods; the meta-classifier model – our best performing model - reclassified 20.9% of individuals deemed high-risk for mortality in logistic regression appropriately as low-to-moderate risk, and 8.2% of deemed low-risk to moderate-to-high risk based consistent with the actual event rates.

**Conclusions:** Machine-learning methods improved the prediction of in-hospital mortality for AMI compared with logistic regression. Machine learning methods enhance the utility of risk models developed using traditional statistical approaches through additional exploration of the relationship between variables and outcomes.

## INTRODUCTION

An assessment of risk of death following an AMI is useful for guiding individual decisions for patients and producing estimates of hospital performance ^1–4^. New analytic approaches may enhance risk prediction with existing data beyond traditional statistical approaches. Previous models have generally neither included non-linear effects for continuous variables nor accounted for complex interactions between variables. With advances in computation and analytics, however, it may be possible to create more accurate models. Specifically, the application of machine learning techniques has the potential to improve upon accuracy in the prediction of in-hospital mortality following AMI ^5–7^

A current gold standard for risk prediction among patients with AMI is derived from the National Cardiovascular Data Registry’s Chest Pain-MI Registry (formerly known as the ACTION Registry, henceforth called the CP-MI Registry), a national quality program from the American College of Cardiology. The CP-MI Registry includes information on AMI hospitalizations at 1,163 hospitals across the United States and includes more than a million patients. Two models for mortality following AMI were recently published using carefully constructed logistic regression models. ^8,9^ The more contemporary model is being used for comparison with alternative approaches. Accordingly, our study evaluated whether the application of machine-learning techniques to data collected in the CP-MI registry could improve upon the prediction of in-hospital AMI mortality over that offered by the well-constructed logistic regression model using these data.

## METHODS

### Chest pain-MI Registry

The CP-MI Registry is a voluntary registry that collects data from participating hospitals on patients admitted with AMI, defined as either ST-elevation myocardial infarction (MI) or non-ST-elevation MI. Patient data are collected through retrospective chart review and submitted to the registry using a standardized data collection tool. Collected data includes patient demographics, presentation information, pre-hospital vital signs, selected laboratory data from the hospital course, procedures, timing of procedures, and select in-hospital outcomes. The NCDR data quality program, which includes audits and feedback, is designed to enhance data completeness and accuracy.^8^

### Patient population

Between January of 2011 and December 2016, a total of 993,905 patients with AMI from 1,128 hospitals were included. We excluded patients transferred to another facility for management (n = 47,308), those missing information on two or more risk factors previously shown to be important for predicting mortality outcomes (n = 191,195). These variables included history of prior MI, history of prior PCI, history of heart failure, history of prior atrial fibrillation, or history of CABG. After exclusion of these patients, 755,402 patients remained for modeling.

### Patient variables and data definitions

Patient variables available to a clinician at the time of presentation were selected for modeling. These variables include patient demographics, history and risk factors, electrocardiogram findings, initial medical presentation information, initial laboratory values, and home medications. The eTable 1 (online supplement) includes a full list of candidate variables. The outcome of our study was death from any cause during hospitalization.

*A priori*, we excluded continuous or categorical variables missing in more than 10% of patients, resulting in 8 candidates (of 14 continuous variables). We calculated three additional continuous variables: body mass index (BMI), creatinine-clearance (using the Cockcroft-Gault equation), and the troponin ratio (value divided by laboratory-specific upper limit of normal). Among categorical variables, 48 of 56 remained after exclusion criteria were applied. The missing rate of the final set of 59 continuous and categorical variables used in modeling was < 1%. For these, we imputed missing values to the mode for categorical variables and median for continuous variables.

### Statistical analysis

Prior to modeling, the cohort was randomly divided 40:40:20 into three samples with equal event rates in each sample for training and validation purposes using the Caret package in R (version 6.0-0.8) To establish a baseline for comparison, we applied the most recent and accepted model based on conventional logistic regression by McNamara et al using the twenty-seven regression variables.^9^

We compared three modeling strategies with logistic regression: 1) logistic regression with lasso regularization (referred to as lasso), 2) gradient descent boosting, and 3) a meta-classification approach combining lasso, a gradient descent boosting, and a neural network. Lasso couples the fitting of a logistic regression model with a cost function that penalizes additional variables, producing a parsimonious set of variables that maximizes predictive capabilities by shrinking the logistic regression coefficients of less important variables to zero. Unlike conventional logistic regression, lasso requires no additional forward or backward selection step to derive a parsimonious variable set ^10^. The lasso was implemented using the glmnet R package (version 2.0-16) ^11^. Gradient descent boosting models make predictions using a series of decision-trees. Unlike logistic regression, tree-based methods can include higher order interactions and account for nonlinear relationships, making them more effective in identifying complex relationships between model variables and outcomes. The method of gradient descent boosting chosen was extreme gradient boosting, or “XGBoost”, And was implemented using the Xgboost R package (version 0.71.1).^12^ Xgboost incorporates a measure of how much model accuracy is improved by the addition of a given variable – a higher gain value implies greater importance in generating a prediction. Finally, the meta-classification approach uses an XGBoost model to combine the outputs of three supervised learning models including lasso, XGBoost, and a neural network (Figure 1). Neural networks are a type of machine learning technique that, like the human brain, connect layers of nodes (neurons), to model an output. The neural network was implemented in Python using the scikit-learn package (version 0.19.1) as an input for metaclassifier approach but was not reported separately.^13^ In the context of our analyses, the lasso and XGBoost models were classified as level 1 classifiers as they were applied to the patient characteristics directly. The metaclassifier was a level 2 classifier as the model was based on the prediction results of the level 1 classifiers.

**Figure 1.**
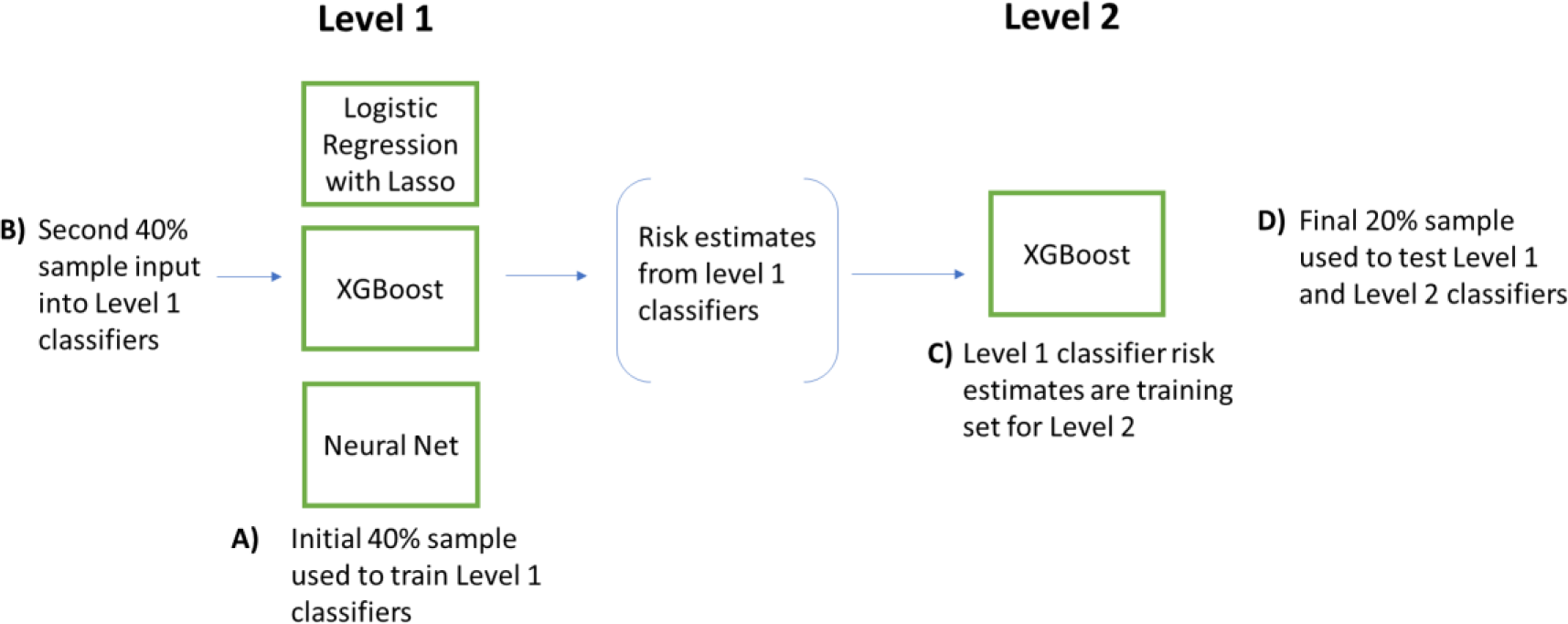
Meta-classifier algorithm design. The level 1 classifiers consist of three independent models each trained on the same initial training sample (A) including logistic regression with lasso, XGBoost, and a neural network. The next training sample (B) is then input into the level 1 classifiers, resulting in three risk estimates for each observation in (B), one from each level 1 model. These three risk estimates are then used to train the level 2 XGBoost classifier (C). A final sample (D) is input into the level 1 classifiers to obtain risk estimates for input into the level 2 classifier. Performance of the level 1 and level 2 classifiers is assessed using this final training set (D).

The computational approach is shown in Figure 1. The first 40% of the data was used to train four methods – logistic regression, lasso, XGBoost, and a neural network. The second 40% of the data were then used as a training set for the level 2 metaclassifier. We validated the various approaches with the remaining 20% of the sample.

In all machine learning models, variables that were imputed were specifically identified and the identifier was included for modeling. None of these flags met statistical significance in any of the models, which suggest that the imputation of these variables was an unlikely source of bias. With the exception of neural network modeling all statistical analysis was done using the open-source R programming language for statistical computing.^14^

The Yale University Institutional Review Board reviewed the study and given the de-identified nature of the data included in the study, waived the requirement for informed consent.

### Performance metrics

Model discrimination was measured using the area under the receiver operator characteristic curve (AUC, or c-statistic) and its 95% confidence intervals ^15^. In addition, the positive predictive value (or precision) and the sensitivity (recall) across all possible risk thresholds for predicting mortality were plotted using the precision-recall curve. The precision-recall curve, unlike the AUC, is unaffected by the number of true negatives. In datasets with a small event rate and therefore a large expected true negative rate, such as the one studied here, the precision-recall curve is well-suited for comparing different models. For both c-statistic and area under the precision-recall curve, values closer to one correspond to more accurate models.

We also calculate the F-score, sensitivity, specificity, positive predictive value (PPV), negative predictive value (NPV). The F-score is the harmonic mean of the sensitivity and PPV at a certain risk threshold, which classifies an individual risk estimate as either a death (if above the threshold) or no death (if below the threshold). Once a risk threshold is set, the number of true positives, false positives, true negatives, and false negatives can be calculated and used to derive sensitivity, specificity, PPV, and NPV. Here, the risk threshold associated with the highest F-score was selected as the overall risk-threshold for the method, and used to determine the sensitivity, specificity, PPV, and NPV for the overall model. The risk threshold is therefore determined using a data-driven approach and optimized for each model.

We also calculated a Brier Score for each model as a measure of model accuracy. It is equal to the “reliability” minus the “resolution”, plus an “error” term, and represents the mean squared error between the observed and predicted risk.^16, 17^ The “reliability” component above is defined as the mean-squared error between the deciles of predicted risk and observed risk – a schematic representation is included in eFigure 1. Notably, since the reliability score measures the error between predicted and observed risk, lower values represent more reliable predictions. The “resolution” component is used to assess the ability of the model to spread patients across a spectrum of risk, which is important for risk stratification of patients. Resolution is calculated as the mean-squared error between deciles of predicted risk and the event rate of the entire cohort. Models with greater resolution have a greater difference between observed mortality rates amongst risk groups and are better performing in that they better spread estimates across a spectrum of risk rather than clustering predictions near the average risk of the population. The “error” term of the Brier Score is based on the event rate in the study cohort and therefore does not vary with different models based on the same cohort. Model calibration was measured using the (a) calibration slope, which was calculated as the regression slope of the observed mortality rates across the deciles of predicted mortality rates, and (b) reliability component of the Brier Score.

Finally, we evaluated whether improvements in risk prediction occurred in low- and high-risk groups, where a change in predicted risk is likely to be clinically significant. For this, we first classified patients in the validation cohort into deciles based on their predicted risk based on logistic regression. Within these deciles, we compared the mean predicted risk based on logistic regression with the mean risk based on each of the three machine learning models and the observed rates of events for individuals within these deciles. We then calculated rates of observed events among individuals reclassified across low, moderate, or high-risk (<1%, 1-5%, >5% risk of mortality, respectively) categories based on logistic regression and one of the machine learning models to identify models that were more likely to capture the risk of patients when there was discordance in the predicted risk based on different modeling strategies.

## RESULTS

### Characteristics of study population

Of the 993,905 individuals with AMI in CP-MI during the study period, 755,402 satisfied our inclusion criteria. The overall in-hospital mortality rate was 4.3%. The derivation cohort consisted of 302,161 patients used to derive the level 1 classifiers, 302,161 to train the meta-classifier model, and the remaining 151,080 patients represented the validation cohort. Table 1 includes characteristics for the derivation and validation cohorts. The mean age at presentation was 64 years, the mean weight was 87 kg, 34% were women, and 85% and 12% were of white and black race, respectively. Of note, 74% of patients had hypertension, 34% had diabetes mellitus, 25% had experienced a prior MI, 13% had a diagnosis of heart failure, and 26% had undergone previous PCI. In addition, 39% presented with an ST-elevation MI, 13% presented in heart failure, 4% in cardiogenic shock, and 4% following a cardiac arrest (**Table 1**).

**Table 1.** Baseline characteristics of derivation and validation cohorts

|  | Derivation (N = 302,161) | Validation (N = 151,080) |
| --- | --- | --- |
| **Demographics** |  |  |
| Age, years | 64 ± 13 | 64 ± 13 |
| Weight, kg | 87 ± 22 | 87 ± 22 |
| Female | 104,291 (34%) | 51,962 (34%) |
| Race white | 256,514 (85%) | 128,112 (85%) |
| Race black | 34,728 (11%) | 17,375 (11%) |
| **Medical History** |  |  |
| History of diabetes mellitus | 102,831 (34%) | 51,631 (34%) |
| History of hypertension | 224,967 (74%) | 112,393 (74%) |
| History of dyslipidemia | 184,628 (61%) | 92,182 (61%) |
| Current/recent smoker | 101,662 (34%) | 50,738 (35%) |
| History of chronic lung disease | 27,216 (9%) | 13,604 (9%) |
| Current dialysis | 7,529 (2%) | 3,894 (3%) |
| History of MI | 75,091 (25%) | 37,560 (25%) |
| History of HF | 37,790 (13%) | 19,072 (13%) |
| Prior PCI | 77,077 (26%) | 38,682 (26%) |
| Prior CABG | 40,252 (13%) | 20,300 (13%) |
| History of atrial fibrillation | 24,798 (8%) | 12,451 (8%) |
| Prior cerebrovascular disease | 36,667 (12%) | 18,342 (12%) |
| Prior peripheral arterial disease | 27,111 (9%) | 13,510 (9%) |
| **Presentation** |  |  |
| Presentation after cardiac arrest | 11,828 (4%) | 13,510 (4%) |
| In cardiogenic shock | 11,437 (4%) | 5,895 (4%) |
| In HF | 38,092 (13%) | 5,780 (13%) |
| Heart rate | 84 ± 23 | 82 ± 24 |
| SBP at presentation | 147 ± 35 | 147 ± 35 |
| **Presentation ECG** |  |  |
| STEMI | 117,078 (39%) | 58,241 (39%) |
| New or presumed new ST-depressions | 33,479 (11%) | 16,766 (11%) |
| New or presumed new T-wave inversions | 22,694 (8%) | 11,344 (8%) |
| Transient ST-segment elevation lasting < 20 min | 3,352 (1%) | 1,674 (1%) |
| **Initial laboratory values** |  |  |
| Troponin Ratio | 3.2 (0.75-13.4) | 3.4 (0.75-13.5) |
| Creatinine, mg/dl | 1.3 ± 1.2 | 1.3 ± 1.2 |
| Creatinine clearance, ml/min | 85 ± 42 | 85 ± 42 |
| Hemoglobin, g/dl | 14 ± 2 | 14 ± 2 |

Values are mean ± SD, n (%), or mean (interquartile range)
HF = heart failure, PCI = percutaneous coronary intervention, CABG = coronary artery bypass graft, SBP = systolic blood pressure, STEMI = ST-segment elevation myocardial infarction

### Comparison of Model Performance

Based upon c-statistics, the machine learning models had significantly superior discrimination compared with the conventional modeling techniques. The area under the precision-recall curves was greatest for the XGBoost and meta-classifier models with values of 0.46 and 0.47, respectively, and smallest for logistic regression model (0.39) (**Figure 2**).

**Figure 2.**
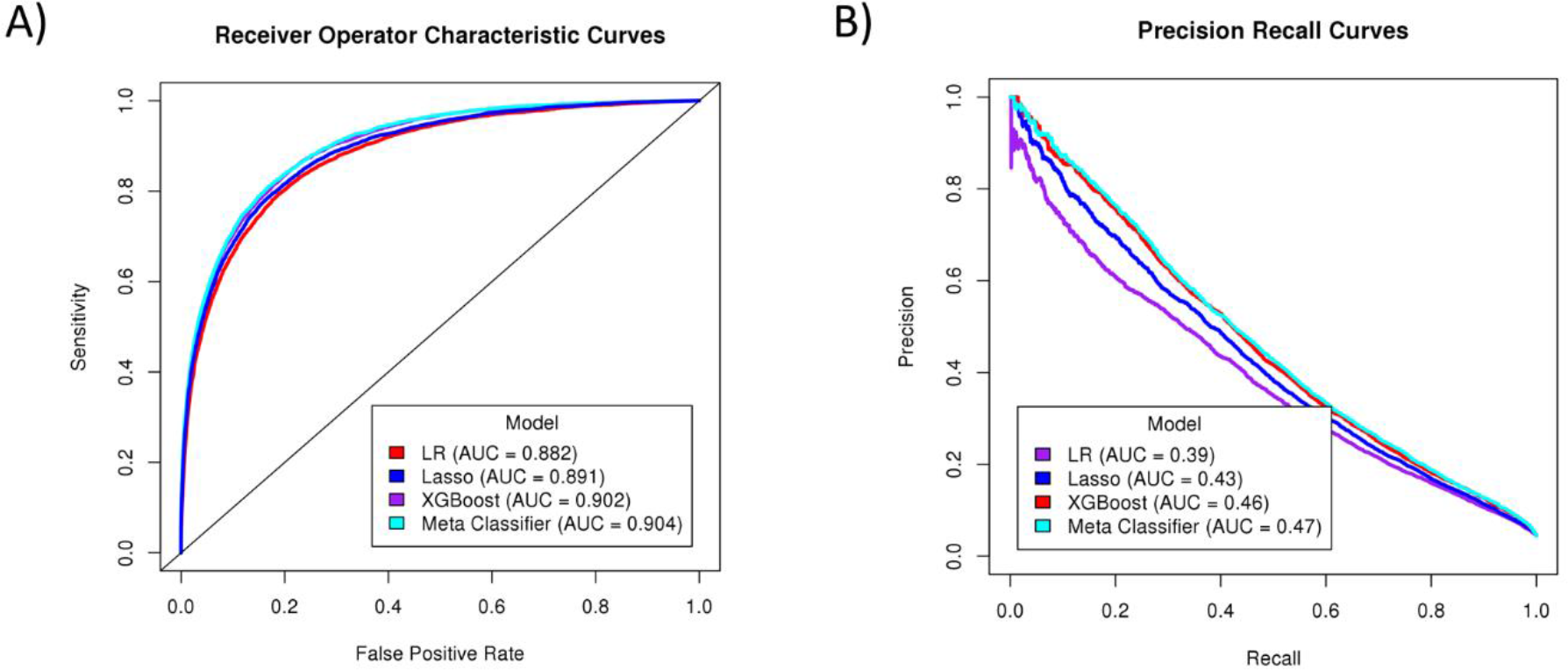
Receiver Operator Characteristic (A) and Precision-recall curves (B) for logistic regression, lasso, XGBoost, and meta-classifier models. (A, left) Receiver Operator Characteristic curves plot the model sensitivity against the false positive rate across a range of all possible risk thresholds for deciding the binary mortality outcome. Curves for baseline logistic regression (LR, purple), lasso (blue), XGBoost (red), and meta-classifier models (cyan) are shown with. The black line shows the performance of an imperfect (random) classifier. Area under the curve (or c-statistic) for each model is shown in the legend. (B, right) Precision-Recall curves for logistic regression (LR, purple), lasso (blue), XGBoost (red), and Meta-classifier models (cyan). Models with precision-recall curves nearest to the top right-hand corner of the graph have the best performance. Area under the curve for each model is shown in the legend.

With respect to calibration slopes, the XGBoost model was 1.006 (95% CI 0.98-1.03), which was a significantly better fit than the logistic regression (0.958 [95% CI 0.94-0.98)]). Neither the lasso model (slope = 0.918, 95% CI 0.90-0.93), or the meta-classifier model (slope = 0.978, 95% CI 0.97-0.98) were superior to the logistic regression model. The components and overall Brier score for the different models are included in Table 2. Models with lower values of “reliability” indicate higher agreement between predicted and observed risk and therefore have better performance. The reliability of the meta-classifier (4.9 × 10^−6^ ± 4.4 × 10^−6^) and XGBoost models (7.3 × 10^−6^ ± 3.3 × 10^−6^) were significantly smaller (therefore, more accurate) compared with the logistic regression model (15.0 × 10^−6^ ± 6.6 × 10^−6^, p < 0.05). The machine learning models had significantly greater resolution – higher range of accurate prediction across the spectrum of risk -than the model based on logistic regression. The meta-classifier had the highest resolution of 7.2 ± 0.2 × 10^−3^, followed by the XGBoost model (7.0 ± 0.2 × 10^−3^), the lasso model (6.6 ± 0.2 × 10^−3^), and finally the logistic regression model (6.1 ± 0.2 × 10^−3^).

**Table 2.**
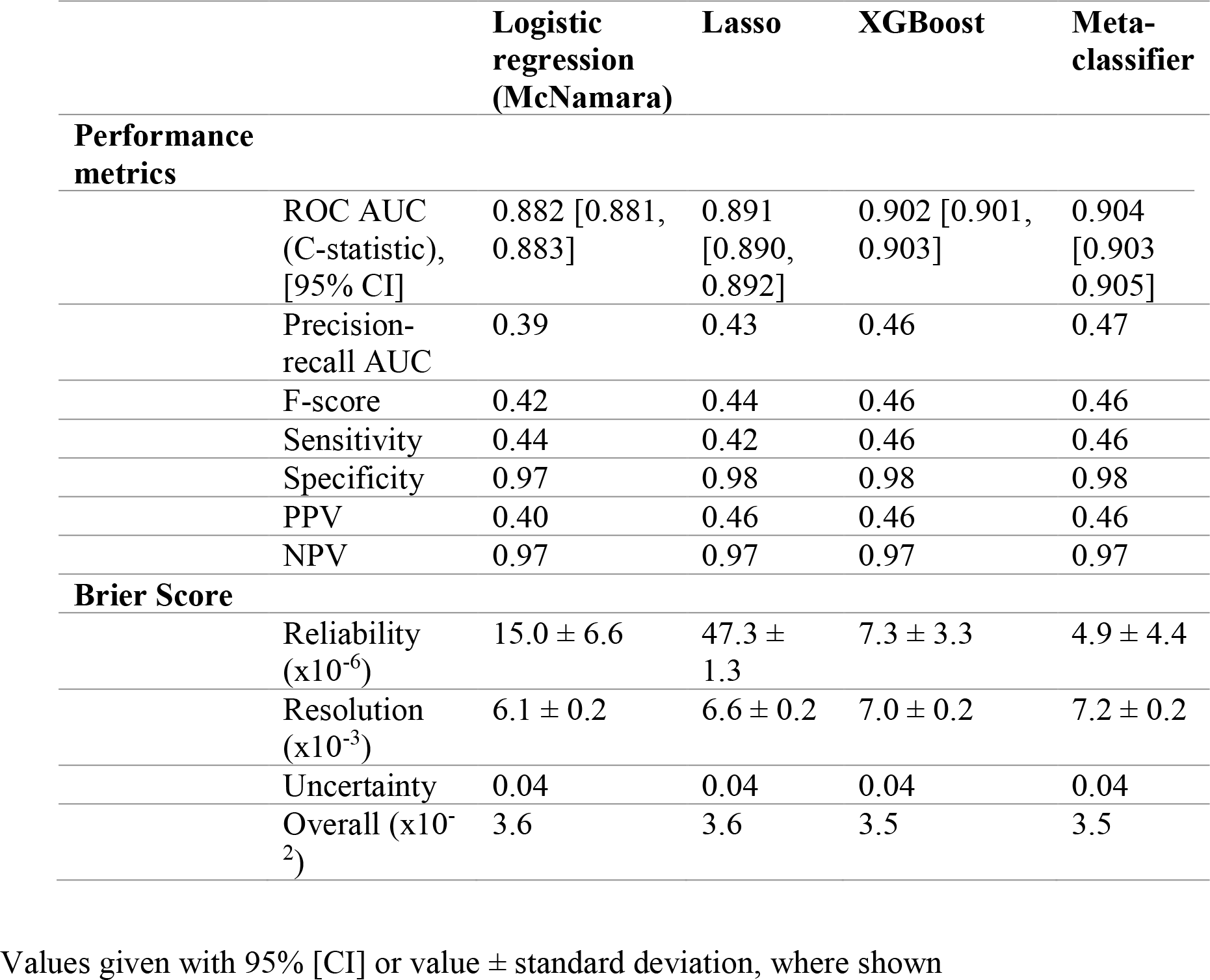
Summary of population-based performance metrics for logistic regression, lasso model, XGBoost model, and meta-classifier models

### Comparison of Individual Risk Prediction

An assessment of risk among those expected to be at the highest and lowest risk based on current prediction algorithms further highlight the improvement offered by machine learning techniques. **Figure 3** presents the risk predicted by all machine learning methods across deciles of risk among the 151,080 patients in the test set based on logistic regression, and the corresponding observed mortality rates in these deciles. Notably, while deciles of predicted risk based on the logistic regression model were consistent with the predicted risk based on the machine learning models across deciles, the mean risk in machine learning models, XGBoost and meta-classifier model more closely approximated the observed risk in these groups (**Figure 3**).

**Figure 3.**
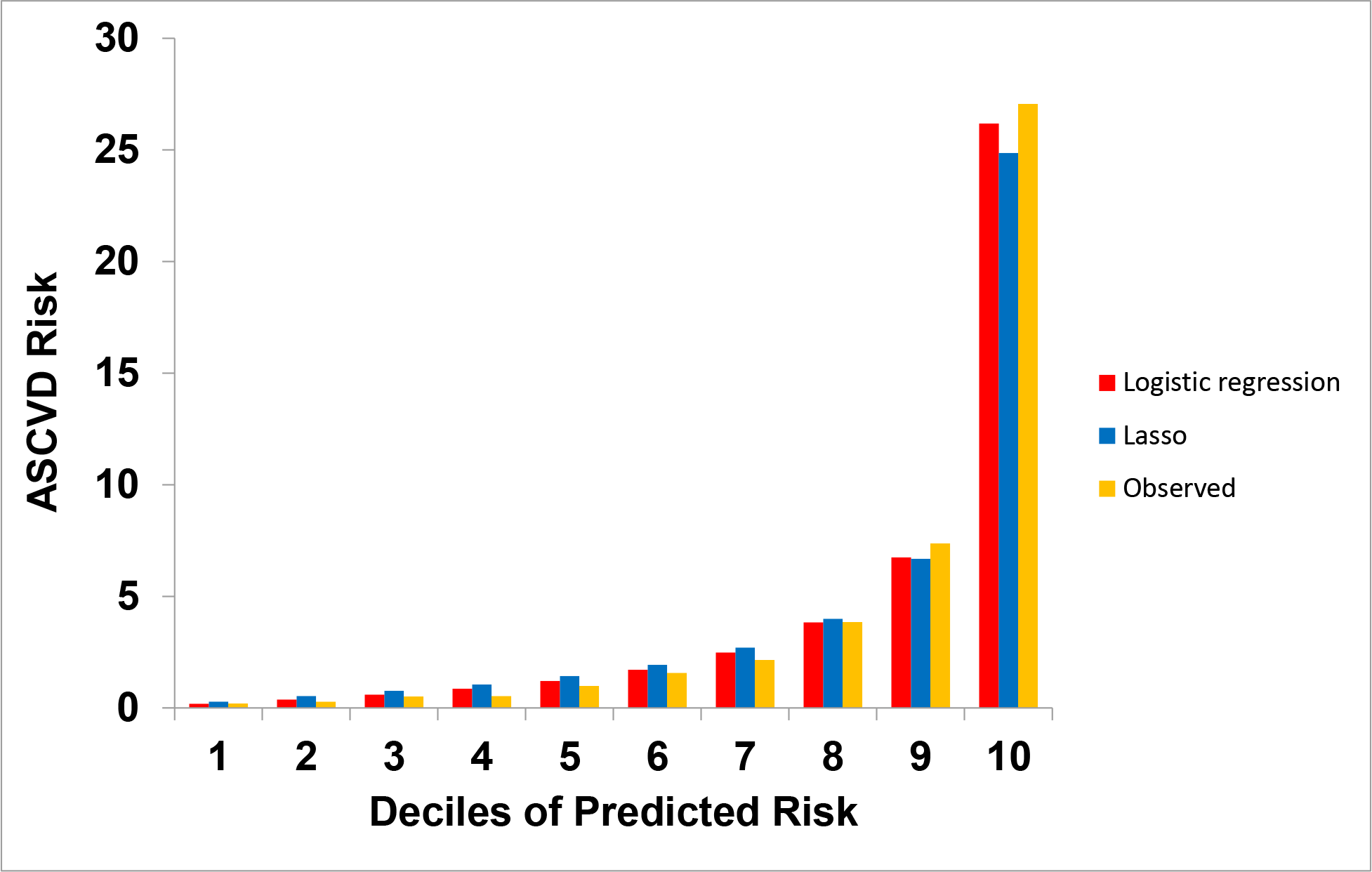
Comparison of individual risk estimates across deciles of risk based on the logistic regression (LR) model. (A) Observed mortality and lasso-predicted risk of mortality

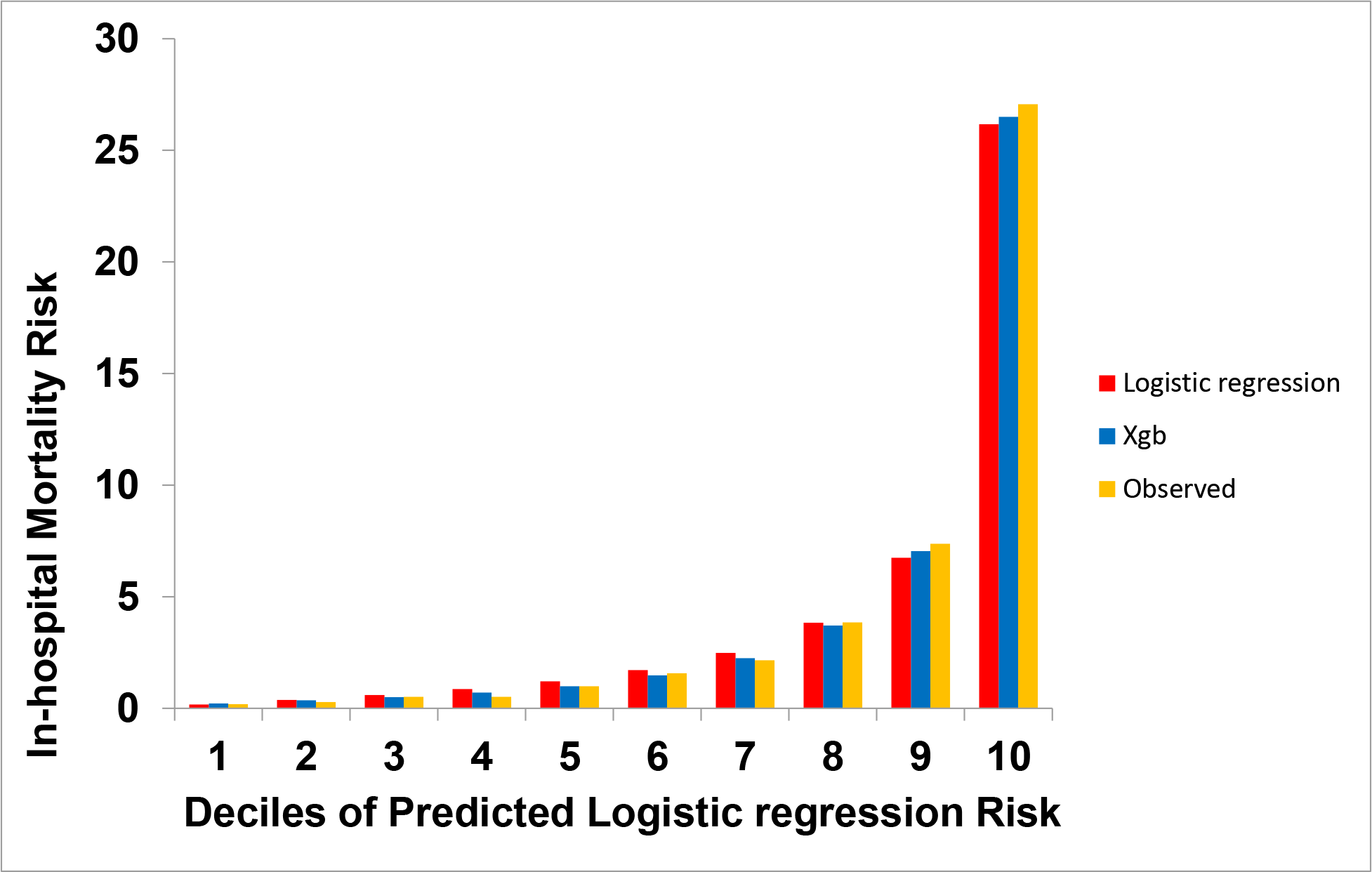
(B) Observed mortality and XGBoost-predicted risk of mortality (Xgb)

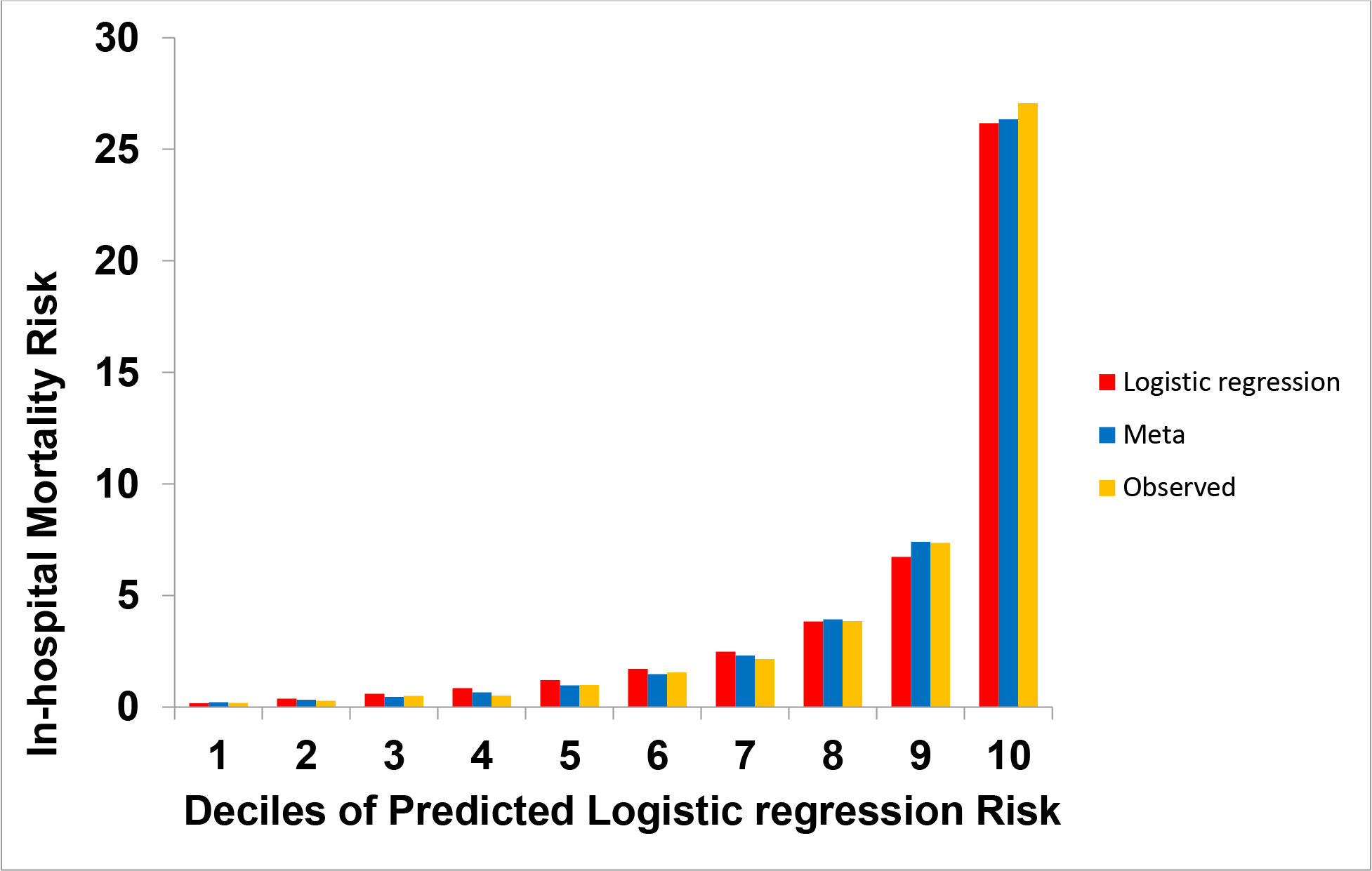
(C) Observed mortality and meta-classifier-predicted risk (meta)

A shift table presented as **Table 3** further illustrates differences between logistic regression and machine learning methods. It shows actual event rates among groups based on their predicted risk across different models (<1%, low risk, 1-5% moderate risk, >5% high risk). These tables highlighted that individuals reclassified by machine learning methods were accurately classified into groups with a rate of events most consistent with their expected risk. For example, among patients predicted to low risk based on the lasso model, and low, moderate or a high risk by logistic regression, there was a negligible difference in the mortality rate among those also predicted to be low-risk by logistic regression (mortality rate, 0.3%) or moderate or high risk (mortality rate, 0.7%), despite logistic regression predicting a risk of mortality of over 1% in the latter group. This suggest minimal improvement in risk prediction over and above the lasso model. In contrast, for patients predicted to be low-risk based on logistic regression, the observed mortality rate among those predicted to be low risk in the lasso model was 0.3% but was 42.9% among those predicted to be high-risk based on the lasso model. Notably, for the latter group of individuals, logistic regression predicted an event rate less than 1%. Similar results are seen across all three machine learning methods (**Table 3**).

**Table 3.**
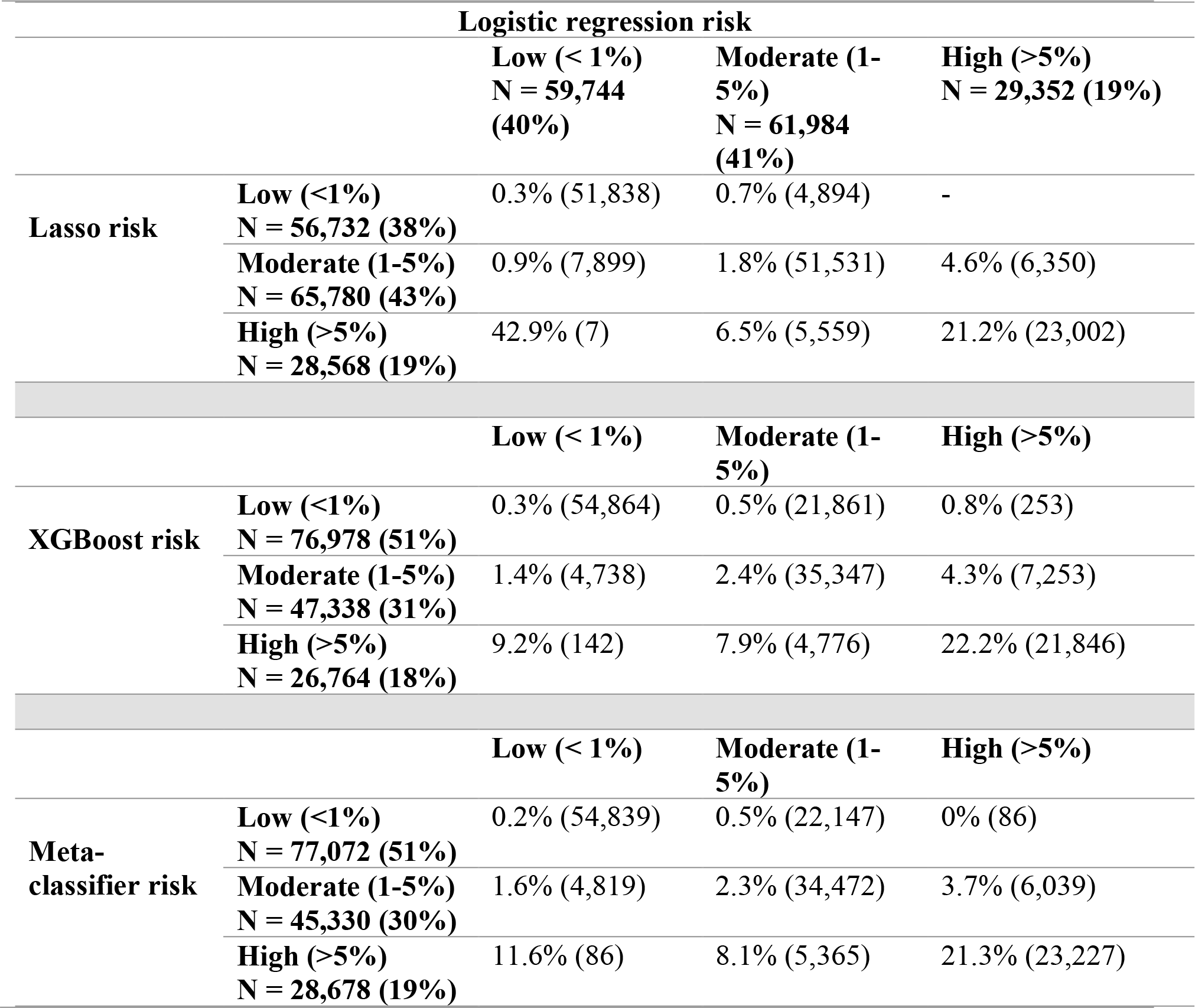
Shift table representing actual observed event rates for pairs of models. Three categories of predicted risk based on the logistic regression are compared against the predicted risk for the same patients using lasso model, XGBoost model, and the meta-classifier (bottom third). **Event rate is reported as a percentage for each cohort, and the cohort size is shown in parentheses.**

Similarly, among patients with a low-risk based meta-classifier model, a moderate/high risk predicted by logistic regression did not reflect an elevated rate of mortality, with an observed mortality rate of <1% across low to high predicted risk by logistic regression (**Table 3**). In contrast, among those with a low predicted logistic regression risk, a low meta-classifier risk was associated with a mortality rate of 0.2%, moderate risk was associated with a mortality of 1.6%, and a high mortality risk by meta-classifier was associated with an observed mortality rate of 11.6%. The techniques also appropriately reclassified patients. Notably, 6125 of 29,352 (or 20.9%) of individuals deemed high-risk in logistic regression were appropriately reclassified as low-to-moderate risk, and 4905 of 59,774 (or 8.2%) deemed low-risk by logistic regression were reclassified as moderate-to-high risk, which was consistent with the actual observed risk.

## DISCUSSION

Our study demonstrates the incremental value of machine learning techniques over traditional logistic models in predicting in-hospital death with AMI in a cohort derived from a US national registry ^9^. Specifically, the new models more precisely identified patient risk across the spectrum of actual risk, with the highest performing machine learning model tested in our study – the meta-classifier – leading to reclassification of 1 in every 5 patients deemed high-risk for mortality in logistic regression accurately as low-to-moderate risk, and 1 in every 12 patients of deemed low-risk to moderate-to-high risk based consistent with the actual observed risk. These observations are particularly notable as the improved performance for models occurs at no additional collection cost and is likely to improve with actual implementation due to the iterative of model performance that occurs in machine learning methods. Moreover, improvements in prediction were achieved those at greatest risk of adverse outcomes, and, therefore, machine learning models have the potential to guide appropriate therapy and outcomes.

Our work presents a proof-of-concept for the improvement in risk-prediction for AMI mortality by applying machine learning techniques. The modeling strategies improve upon the application of logistic regression models that are limited in their performance by the characteristics of the data as well as correlations between variables available for risk-prediction.^5–7^ In addition, some models have traditionally relied on literature review or expert opinion for variable selection used in predictive modeling, which can further lead to loss of information about potential predictors and relationships.

Our findings have several important implications. First, through more accurate and precise prediction of risk across the spectrum of individual mortality risk, these advanced modeling techniques have the potential to better calibrate the intensity of treatment to risk of adverse outcomes and to improve the communication of the risk for adverse outcomes to patients and their families. An accurate assessment of risk is critical for addressing goals of care and treatment strategies. While the overall improvement in prediction accuracy of the machine learning models over logistic regression was modest, through greater predictive range, these improvements occurred in critical areas by accurately re-classifying individuals at low and high-risk to categories more accurately reflecting their risk. Notably, nearly 5000 individuals were expected to be at low-risk for in-hospital death based on the currently accepted risk-assessment, guiding both their in-hospital treatment and communicating their risk to patients for shared decision-making, but were accurately noted to be at a much higher risk based on the machine learning models. Second, the improvement in prediction of risk of adverse outcomes based on presentation characteristics should be able to improve the profiling of hospital performance through better adjustment for baseline patient risk. The study offers insights into the future research on the nature of improvements to risk-prediction offered through application of machine learning techniques to the study of healthcare delivery. The advantage of a greater reliance on these techniques will also manifest with their clinical application. These models offer an opportunity to continually improve risk-prediction in real time, especially as predictors and their relationship to each other evolve over time. This is an important feature of machine learning strategies that makes them more adept at clinical applications over other risk-prediction algorithms relying on static models for their derivation, or requiring manual recalibration.

Our study has several limitations. First, while the CP-MI registry captures granular clinical data on AMI patients, relevant information such as duration of comorbidities, control of chronic diseases (besides diabetes) were not captured and are, therefore, not included in our assessment. Second, machine learning methods assess complex relationship between predictors and are not accompanied by usual associations for individual variables and outcomes based on clinical judgement or empirical evidence of associations. Notably, however, the models are based on sound mathematical principles and are easily replicated in these data. The study, also does not identify whether the excess risk identified for patients using these models is actually reversible through improved care, and needs dedicated investigation in the future. Finally, the study is generalizable to the data in the NCDR CP-MI Registry and may not apply to patients not included or hospitals not participating in the registry. However, since the data included in the modelling strategy are collected as a part of routine clinical care at participating hospitals, other hospitals collecting similar data could likely apply these modeling strategies.

In conclusion, we developed machine learning models that outperformed carefully designed risk-prediction models based on traditional methodology in the prediction of death following an AMI. These models offer greater resolution of risk without collection of additional data and can better clarify individual risk for adverse outcomes and more accurately compare hospital performance for AMI care.

## Online Supplement

**eTable 1:**
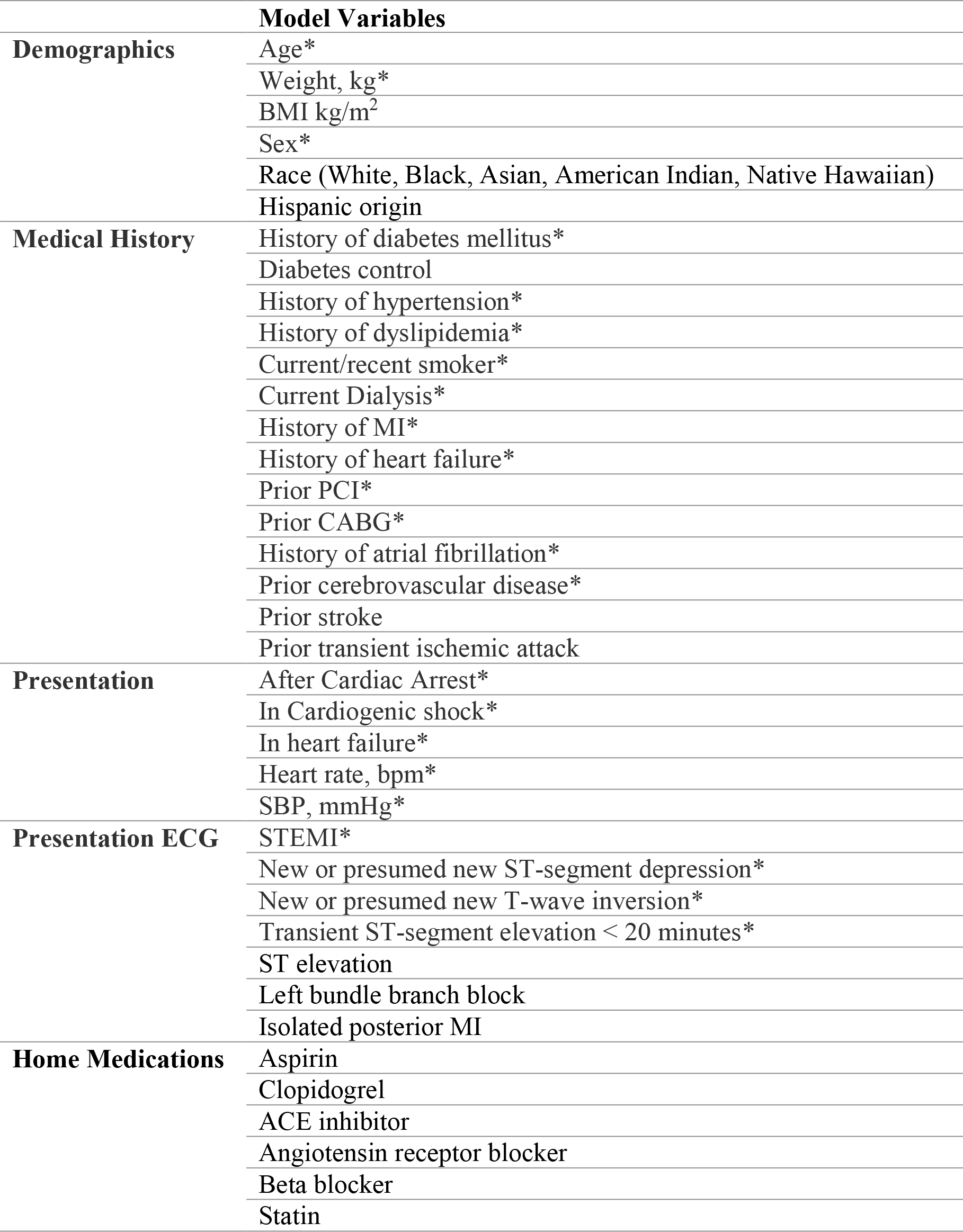
List of patient variables used in modeling. * denotes model variables used in McNamara et al. study.^9^ Creatinine clearance calculated via Cockgroft-Gault equation.

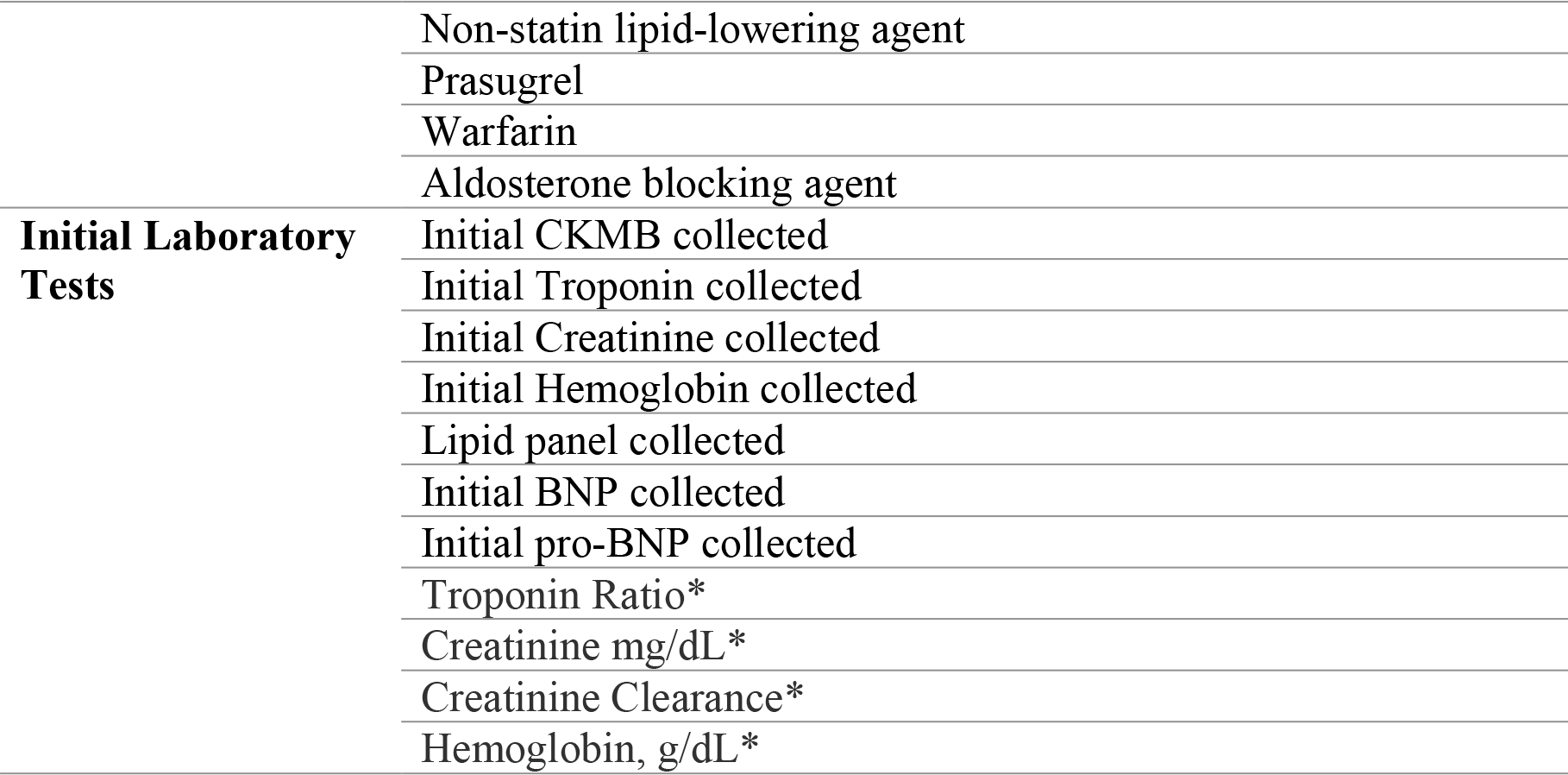

**eTable 2.** Importance of level 1 models to meta-classifier

| **Level 1 Model** | **Importance (%)** |
| --- | --- |
| **XGBoost** | 65.3 |
| **Neural network** | 34.6 |
| **Logistic regression with lasso** | 0.1 |

Importance describes the relative contribution of each of the Level 1 models to the output of the meta-classifier as measured by the XGBoost algorithm. Here, the Level 1 XGBoost model is the major contributor to the meta-classifier model followed by the neural network.

**eFigure 1.**
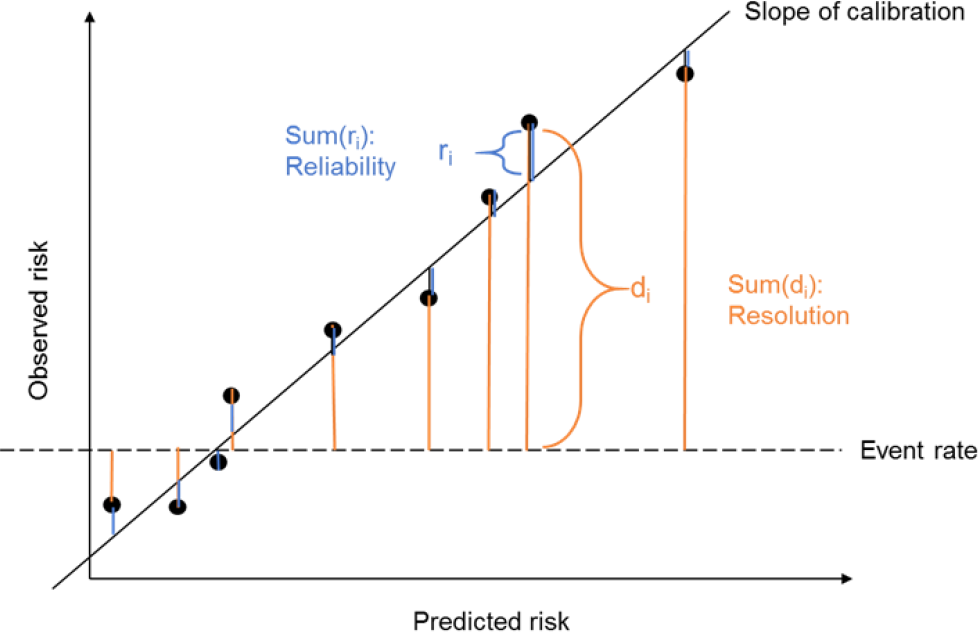
Derivation of Brier Score components based on calibration curve. In the figure, each point represents the predicted versus observed risk at a given decile of risk. Reliability is the sum of the mean-squared error between the deciles of predicted risk and observed risk, and Resolution is the mean-squared error between deciles of predicted risk and the event rate of the entire cohort

**eFigure 2.**
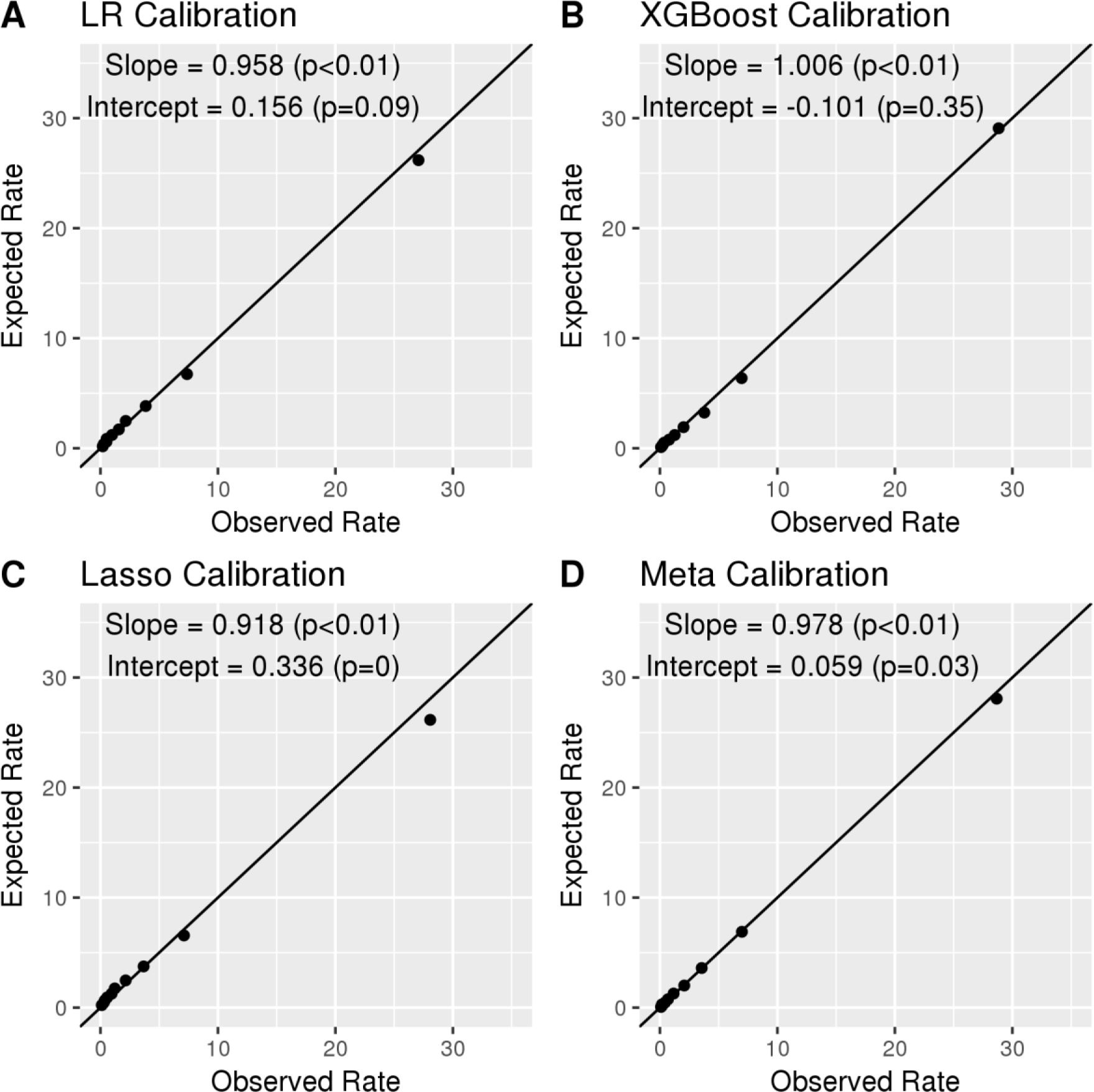
Calibration curves for logistic regression (LR, A), XGBoost (B), Lasso (C) and Meta-Classifier (d) models for validation cohort predictions. Slope and intercept of the linear fit to each calibration plot are given in the top-left of each pane with corresponding p-values for each coefficient, where a p-value < 0.05 for the slope coefficient indicates that the slope is non-zero and that there is a significant association between the observed and expected rate, and a p-value < 0.05 for the intercept coefficient indicates that the intercept is non-zero.

